# A general model of multivalent binding with ligands of heterotypic subunits and multiple surface receptors

**DOI:** 10.1101/2021.03.10.434776

**Authors:** Zhixin Cyrillus Tan, Aaron S. Meyer

**Author notes:** Corresponding author *Email address:* (Aaron S. Meyer).

## Abstract

Multivalent cell surface receptor binding is a ubiquitous biological phenomenon with functional and therapeutic significance. Predicting the amount of ligand binding for a cell remains an important question in computational biology as it can provide great insight into cell-to-cell communication and rational drug design toward specific targets. In this study, we extend a mechanistic, two-step multivalent binding model. This model predicts the behavior of a mixture of different multivalent ligand complexes binding to cells expressing various types of receptors. It accounts for the combinatorially large number of interactions between multiple ligands and receptors, optionally allowing a mixture of complexes with different valencies and complexes that contain heterogeneous ligand units. We derive the macroscopic predictions and demonstrate how this model enables large-scale predictions on mixture binding and the binding space of a ligand. This model thus provides an elegant and computationally efficient framework for analyzing multivalent binding.

## 1. Introduction

Binding to extracellular ligands is among the most fundamental and universal activities of a cell. Many important biological activities, and cell-to-cell communication in particular, are based on recognizing extracellular molecules via specific surface receptors. For example, multivalent ligands are common extracellular factors in the immune system [1], and computational models have been applied to study IgE-FcεRI [2], MHC-T cell receptor [3], and IgG-FcγR interactions [4]. However, these models are specific to their biological applications, limited to a single homogenous ligand and receptor [5], or fail to scale with valency [6].

Multivalent binding to various receptors on a cell can be accounted for by the kinetics of individual association reactions between each monomer-receptor pair. However, when the complexes contain multiple ligand monomers of either the same or different kinds, and when there is a mixture of complexes with either the same or different valencies, the system becomes complicated: different binding orders of units on a complex creates a combinatorially large amount of possible reactions, and the competition among different kinds of ligands and complexes impedes intuitive understanding. In this case, enumerating all binding configurations and reactions become impractical.

In this study, we extend a simple two-step, multivalent binding model to cases involving multiple receptors and ligand subunits [7, 8, 5, 9, 3]. By harnessing the power of combinatorics via applying the multinomial theorem and focusing on macrostates, we can predict the amount of binding for each ligand and receptor at equilibrium. We derive macroscopic quantities for both specifically arranged and randomly assorted complexes, and demonstrate how this model enables large-scale predictions on mixture binding and the binding space of a ligand.

Our model provides both generality and computational efficiency, allowing large-scale predictions such as characterizing synergism of using a mixture of ligands and depicting the general binding behavior of a compound. The compactness and elegance of the formulae enable both analytical and numerical analyses, in turn allowing for the construction of higher-level computational tools. We expect this binding model will be widely applicable to many biological contexts.

## 2. Preliminaries

### 2.1. Vector and matrix notation

In this work, we denote a vector in boldface letter and its entry in the same letter but with subscript and not in boldface, e.g. **C** = [*C*_1_, *C*_2_, …, *C*_*n*_]. The sum of elements for a vector is denoted as 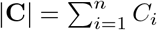.

For any matrix (*A*_*ij*_) of size *m* × *n*, we denote the vector formed by its *i*-th row as **A**_**i**•_ = [*A*_*i*1_, *A*_*i*2_, *…, A*_*in*_], and the vector formed by its *j*-th column as **A**_•**j**_ = [*A*_1*j*_, *A*_2*j*_, *…, A*_*mj*_]. The row sums of matrix (*A*_*ij*_), therefore, can be written as |**A**_**1**•_|, |**A**_**2**•_|, *…*, |**A**_**m**•_|, and column sums |**A**_•**1**_|, |**A**_•**2**_|, *…*, |**A**_•**n**_|.

In this work, multinomial coefficients such as *n* choose *k*_1_, *k*_2_,…, *k*_*n*_ will be written as

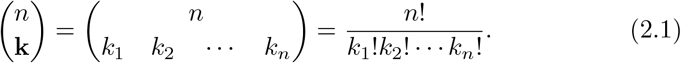

The implicit assumption here is that |**k**| = *n*, and each *k*_*i*_ ∈ ℕ.

### 2.2. Some useful theorems in combinatorics

From the binomial theorem, we know that

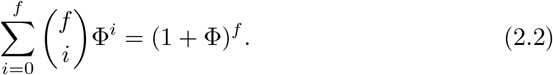

Differentiating both sides by Φ, we get

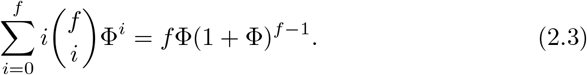

We can derive similar property from the multinomial theorem. Assume the elements of a nonnegative integer vector **q** add up to *f*, or |**q**| = *f*. Given another nonnegative vector ***φ*** with sum of elements |***φ***|, we have

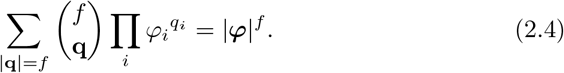

Differentiate both sides by *φ*_*m*_ where *φ*_*m*_ can be any entry of ***φ***, we have

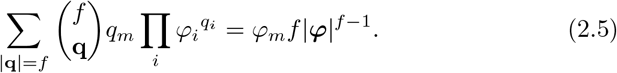

We can multiply two independent multinomial theorem equations together, too. Let **u** and **v** be two nonnegative integer vectors, **a** and **b** be two nonnegative vectors, and |**u**| = *m*, |**v**| = *n*, we have

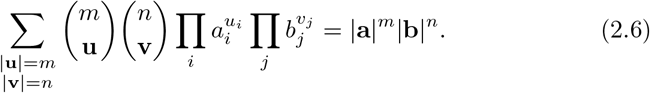

Throughout this work, we consolidate multiple summation symbols into one. In this case, we use Σ_|**u**| =*m*,|**v**| =*n*_ as a shorthand for Σ_|**u**| =*m*_ Σ_|**v**| =*n*_. From Equation 2.5, we can derive the sum of a linear combination of two exponents from each multinomial term as

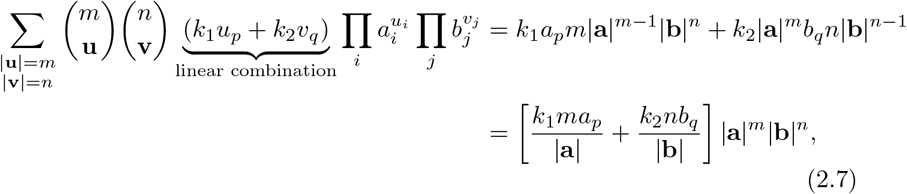

where *k*_1_ and *k*_2_ can be any constant.

We can extend this to the product of *N* multinomial equations. Let **q**_**1**_, *…*, **q**_**N**_ be *N* nonnegative integer vectors, each with |**q**_**i**_| = *θ*_*i*_, and ***ψ***_**1**_, *…*, ***ψ***_***N***_ be *N* nonnegative vectors. The sum of any linear combination of exponent terms 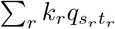, where *k*_*r*_’s can be any constant, and each 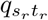 is the *t*_*r*_-th element of 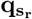, can be calculated as

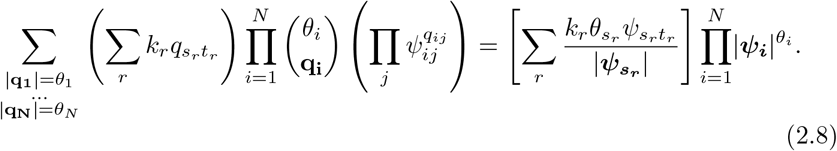

## 3. Model setup

### 3.1. Parameters and notations

In this study, we investigate the binding between multivalent ligand complexes and a cell expressing various surface receptors. As shown in Figure 1, we consider *N*_*L*_ types of distinct monomer ligands, namely 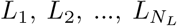 and *N*_*R*_ types of distinct receptors expressed on a cell, namely 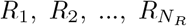.

**Figure 1:**
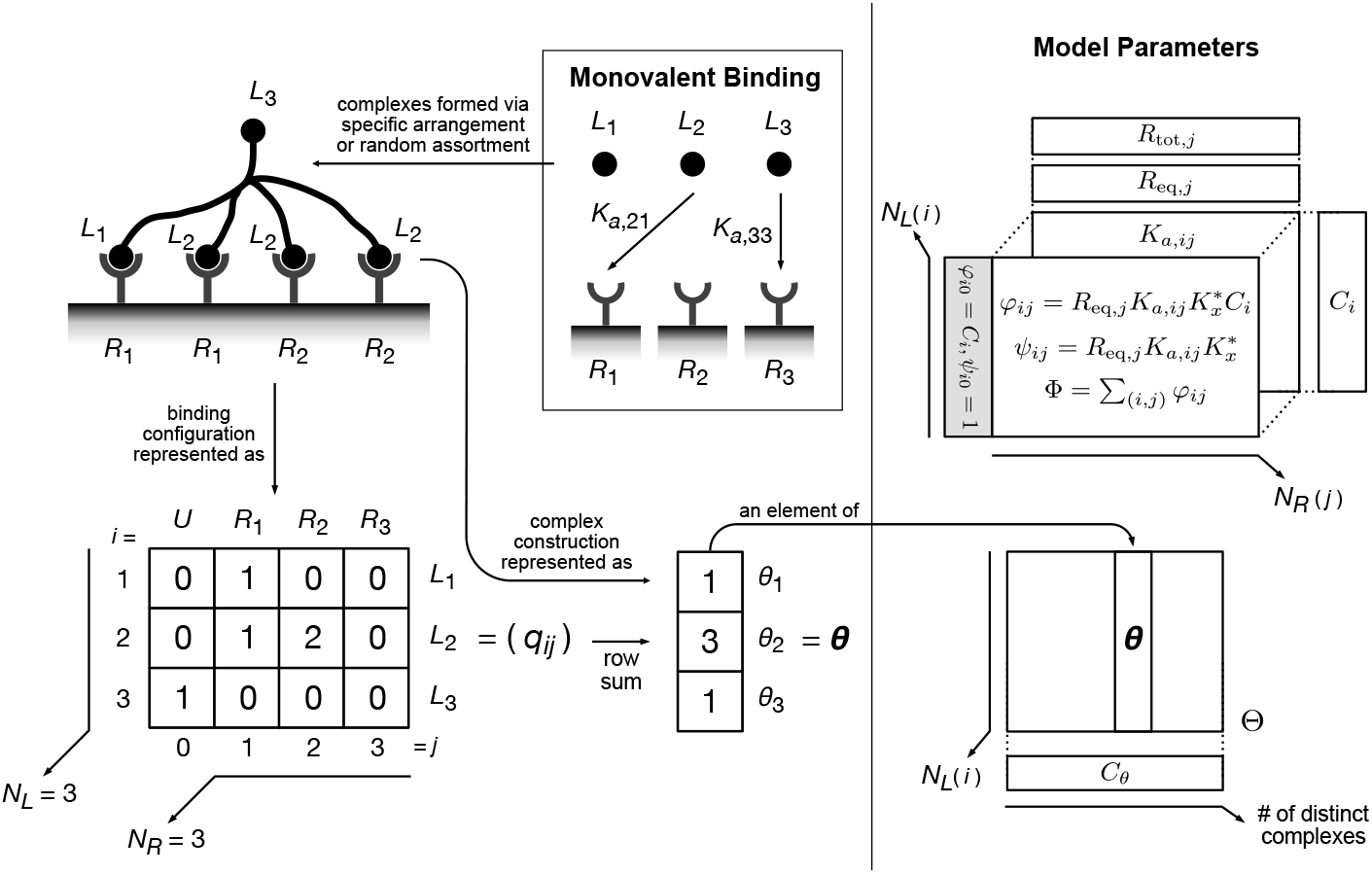
General setup of the model. In this study, we investigate the binding behavior of complexes formed by monomer ligands in either a specific arrangement or by random assortment. We propose that the binding configuration between a complex and several receptors on a cell can be described as a matrix (*q*_*ij*_). The construction of a complex can be written as a vector ***θ***. The figure shows the dimensions of the model’s parameters: *C*_*i*_, the monomer compositions, are in a vector of length *N*_*L*_; *R*_tot,*j*_ and *R*_eq,*j*_, the receptor expression and equilibrium levels are in vectors of length *N*_*R*_; the binding affinities, *K*_*a,ij*_, are in a matrix of dimension *N*_*L*_ × *N*_*R*_; *φ*_*ij*_ and *ψ*_*ij*_ are in the matrices of dimension *N*_*L*_ × (*N*_*R*_ + 1). Θ is a set of all possible ***θ***’s, with *C*_***θ***_ as each of their compositions. Each ***θ*** is a vector of length *N*_*L*_, and *C*_***θ***_ should be in a vector of the same size as Θ.

The monovalent binding association constant between *L*_*i*_ and *R*_*j*_ is defined as *K*_*a,ij*_. A ligand complex consists of one or several monomer ligands, and each of them can bind to a receptor independently. Its construction can be described by a vector 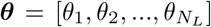, where each entry *θ*_*i*_ represents how many *L*_*i*_ this complex contains. The sum of elements of vector ***θ***, |***θ*** |, is *f*, the valency of this complex.

The binding configuration at equilibrium between an individual complex and a cell expressing various receptors can be described as a matrix (*q*_*ij*_) with *N*_*L*_ rows and (*N*_*R*_ + 1) columns. For example, the complex bound as shown on the top left corner in Figure 1 can be described as the matrix below it. *q*_*ij*_ represents the number of *L*_*i*_ to *R*_*j*_ binding, and *q*_*i*0_, the entry on the 0-th column, is the number of unbound *L*_*i*_ on that complex in this configuration. This matrix can be unrolled into a vector form 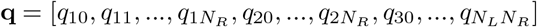 of length *N*_*L*_(*N*_*R*_ + 1). Note that this binding configuration matrix (*q*_*ij*_) only records how many *L*_*i*_-to-*R*_*j*_ pairs are formed, regardless of which exact ligand on the complex binds. For example, in Figure 1, swapping the two *L*_2_’s binding to *R*_2_’s will give us the same configuration matrix. Therefore, we will need to account for this combinatorial factor when applying the law of mass action.

We know from the conservation of mass that for this complex, 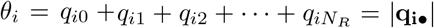 must hold for all *i*’s. Mathematically, vector ***θ*** is the row sums of matrix (*q*_*ij*_). The corresponding ***θ*** of a binding configuration **q, *θ***(**q**), written in the format of a function of **q**, can be determined by this relationship. Also, the sum of elements in **q**, |**q**| = *f*, will be the valency.

The concentration of complexes in the solution is *L*_0_ (not to be confused with *L*_*i*_, the name of ligands, when *i* = 1, 2, *…, N*_*L*_). It is the concentration of all ligands at the equilibrium state. It is approximately the same as the initial ligand concentration if the amount of ligands is much greater than that of the receptors and binding event does not significantly deplete the ligand concentration.

On the receptor side, *R*_tot,*i*_ is the total number of *R*_*i*_ expressed on the cell surface. This usually can be measured experimentally. *R*_eq,*i*_ is the number of unbound *R*_*i*_ on a cell at the equilibrium state during the ligand complex-receptor interaction, and usually must be calculated from *R*_tot,*i*_ as we will explain later.

The binding of a ligand complex, a large molecule, is complicated. To simplify the matter, we will need to make some key thermodynamic assumptions.

In this model, we make two assumptions on the binding dynamics:

1. The initial binding of *L*_*i*_ on a free (unbound) complex to a surface receptor *R*_*j*_ has the same affinity (association constant, *K*_*a,ij*_) as that of a monomer ligand *L*_*i*_;
2. In order for the detailed balance to hold, the association constant of any subsequent binding event on the surface of a cell after the initial interaction must be proportional to their corresponding monovalent affinity. We assume the subsequent binding affinity in multivalent interactions between *L*_*i*_ and *R*_*j*_ to be 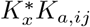.

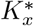 is a term coined as the crosslinking constant. It captures the difference between free and multivalent ligand-receptor binding, including but not limited to steric effects and local receptor clustering [10]. In practice this term is often fit to apply this model to a specific biological context.

We create two last variables that will help to simplify our equations. For all *i* in *{*1, 2, …, *N*_*L*_*}*, we define 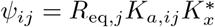 and 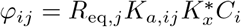 where *j* = *{*1, 2, …, *N*_*R*_*}*, and we define *ψ*_*i*0_ = 1, *φ*_*i*0_ = *C*_*i*_. Therefore, *φ*_*ij*_ = *ψ*_*ij*_*C*_*i*_ holds for all *i* and *j*. Then we define the sum of this new matrix (*φ*_*ij*_) as 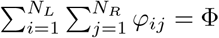, and 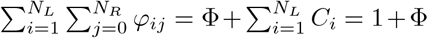. The rationale of these definitions will become clear in future sections.

### 3.2. The amount of a specific binding configuration

With the definitions of our model we can now derive the amount of complexes bound with the configuration described as **q** on a cell at equilibrium, *v*_**q**_. We know that the composition of any complex can be described by a vector ***θ*** of length *N*_*L*_, where each entry *θ*_*i*_ represents the number of monomers *L*_*i*_ within the complex. We can enumerate all possible binding configurations of ***θ*** complex by filling the matrix (*q*_*ij*_) with any nonnegative integer values so long as its row sums equal ***θ***. Conversely, for a certain binding configuration, **q**, the construction of the complex involved must be its row sum, ***θ***(**q**), and the concentration of this complex is *L*_0_*C*_***θ***(**q**)_. If the corresponding complex ***θ***(**q**) does not exist in the solution, we set *C*_***θ***(**q**)_ = 0. Since ***θ***(**q**) is defined only by the ligand concentration at equilibrium, it will remain *L*_0_*C*_***θ***(**q**)_.

#### Initial binding

We start with the initial binding reaction of a complex, *L*_*i*_-to-*R*_*j*_. As shown in Figure 2, the reactants are the free complexes and the free receptors *R*_*j*_ (in this case *R*_2_), and the product are the *L*_*i*_-to-*R*_*j*_ (in this case *L*_2_-*R*_2_) monovalently bound complexes **q**_(**1**)_. We denote the amount of this new complex as 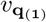. The concentration of free complexes is 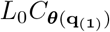. The equilibrium constant for this reaction is *K*_*a,ij*_. Therefore, we have

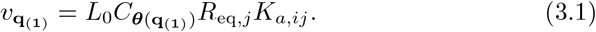

While the binding configuration of **q**_(**1**)_ can be described by **q**_**a**_, the total amount of complexes that bind as described as **q**_**a**_ may not be the same as 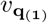, since **q**_**a**_ does not consider the number of ways this binding *L*_*i*_ can be chosen. An equivalent explanation is that, **q**_(**1**)_ is only one possible microstate to achieve the **q**_**a**_ configuration, and we need to count the total number of possible microstates for **q**_**a**_. Accounting for this statistical factor, we have

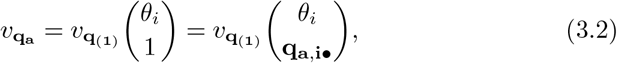

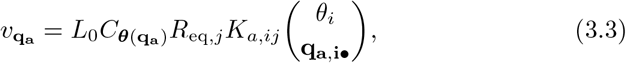

since ***θ***(**q**_(**1**)_) = ***θ***(**q**_**a**_). **q**_**a**,**i**•_ is a vector formed by the *i*-th row of **q**_**a**_. For example, in Figure 2, **q**_**a,2**,•_ = [2, 0, 1, 0]. Conceptually, 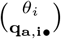 can be understood as the number of ways to split *θ*_*i*_ *L*_*i*_’s into *q*_*i*0_ of unbound, *q*_*i*1_ of *R*_1_-bound, *q*_*i*2_ of *R*_2_-bound, …, and *qiN*_*R*_ of 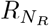 -bound units. In the initial binding reaction, only *q*_*i*0_ and *q*_*ij*_ will be nonzero, with *q*_*i*0_ = *θ*_*i*_ − 1 and *q*_*ij*_ = 1, so it is effectively the same as 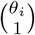. However, the multinomial coefficient expression can be generalized to subsequent binding steps.

**Figure 2:**
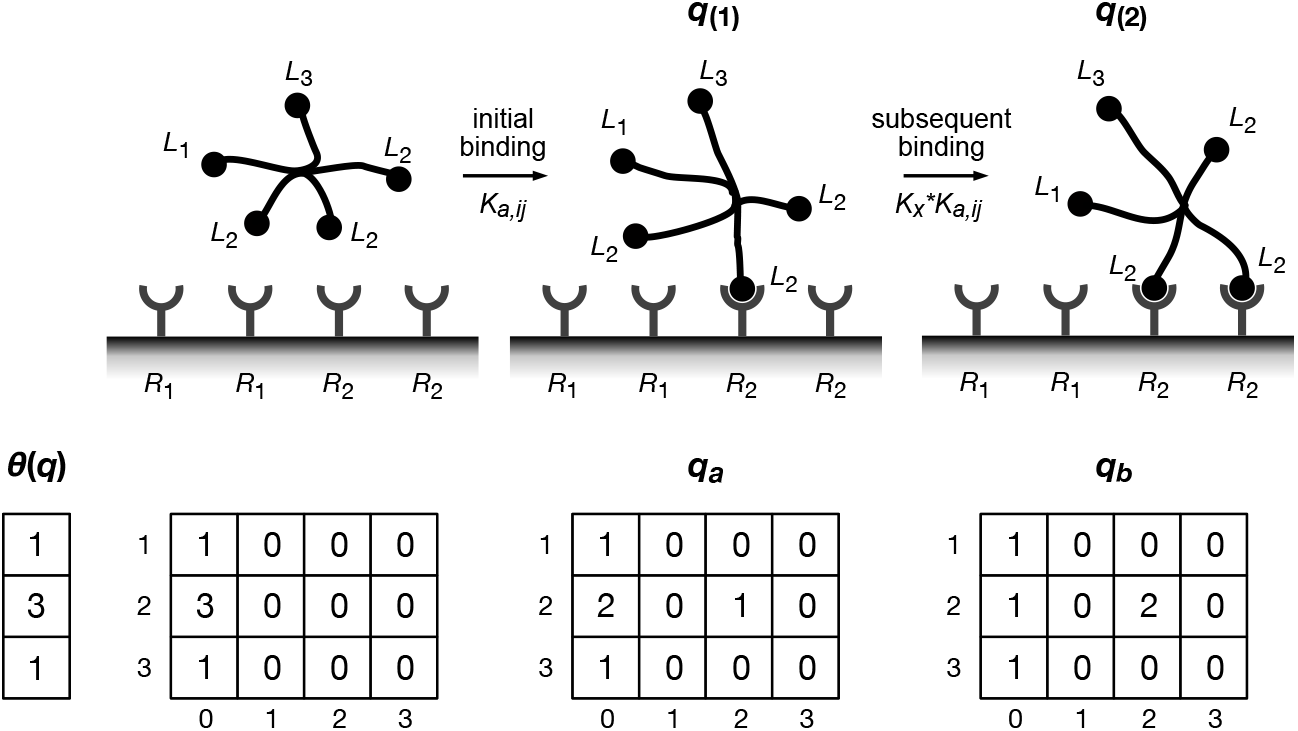
The scheme of cell-complex binding step by step. We assume the initial binding event has the same affinity as monomer binding, *K*_*a,ij*_, while subsequent binding has an association constant scaled by 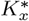, the crosslinking constant. Each binding configuration scheme above can be described by the **q** right below, if we ignore the statistical factors. ***θ***(**q**) is the construction of the complex and can be implied from **q**.

#### Subsequent binding

For a subsequent binding between *L*_*i*_ and *R*_*j*_ (*i* and *j* are not necessarily the same as in initial binding), we have the reactants as a bound complex, **q**_(**1**)_, and a free receptor *R*_*j*_ (in the case shown by Figure 2, *R*_2_), while the product is another bound complex, **q**_(**2**)_. The equilibrium constant is 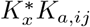, then

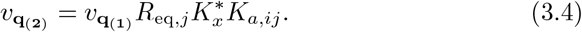

To account for the statistical factors of 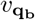, we have 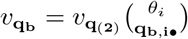. For example, in Figure 2, **q**_**b**,**2**•_ = [1, 0, 2, 0]. Putting these together, we have

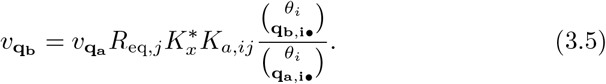

By recursion, we can solve *v*_**q**_ for any **q** from these equations. It is

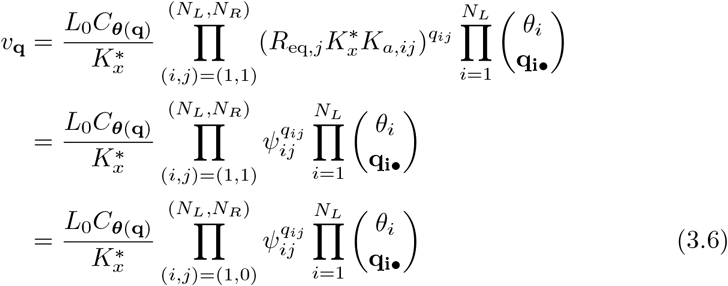

if we define 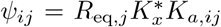 for *j* = 1, 2, *…, N*_*R*_ and *ψ*_*i*0_ = 1 for all *i*. 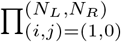 is a shorthand for 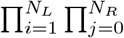. In the next section, we will use this formula repeatedly.

Notice that this equation is only for surface-bound complexes, not suitable for calculating the concentration of unbound **q**, when every nonzero values are on its 0-th column. The concentration of unbound ligands should always be *L*_0_*C*_***θ***(**q**)_. However, for algebraic convenience, we allow such definition but only to subtract them later, and will name it *v*_0,eq_ which equals 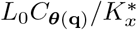.

## 4. Macroscopic equilibrium predictions

From here we will investigate the macroscopic properties of binding, such as the total amount of ligand bound and receptor bound on a cell surface at equilibrium. We consider two different ways complexes in the solution can be formed. First, complexes may come in a specific arrangement. In this case, the structure and exact concentration for each complex are designed and known. Alternatively, ligand monomers of known proportion can congregate into complexes with a fixed valency *f*. Through random assortment, any combination of *f* monomer ligands can form a complex, and their concentration will follow a multinomial distribution. We will explore these two cases separately.

### 4.1. Complexes formed in a specific arrangement

When complexes are specifically arranged, the structure and proportion of each kind are well-defined. To formulate this mathematically, we assume that we have various kinds of complexes, and each of them can be described by a vector ***θ*** of length *N*_*L*_, with each entry *θ*_*i*_ as the number of *L*_*i*_ in this complex. The valency of each complex may be different, and for complex ***θ*** its valency is |***θ***|. The proportion of ***θ*** among all complexes is defined as *C*_***θ***_, and the concentration of each ***θ*** complex will be *L*_0_*C*_***θ***_. For example, if we create a mixture of 20% of bivalent *L*_1_ and 80% of bispecific *L*_1_ − *L*_2_, then ***θ***_**1**_ = [2, 0], ***θ***_**2**_ = [1, 1], 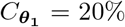, and 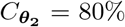. If the mixture solution has a total concentration of 10 nM, then the concentration of ***θ***_**1**_ is 2 nM, and the concentration of ***θ***_**2**_ is 8 nM.

We further conceptualize that Θ is a set of all existing ***θ***’s. By this setting, we should have Σ_***θ∈***Θ_ *C*_***θ***_ = 1. These complexes will bind in various configurations which can all be described as a **q**. We define *Q* as a set of all possible **q**’s, and we borrow the notation **q** ⊆ ***θ*** to indicate any binding configuration **q** that can be achieved by complex ***θ***. This is equivalent to |**q**_**i**•_| = *θ*_*i*_ for all *i*, or ***θ*** is the row sum of (*q*_*ij*_).

#### Solve the amount of free receptors

A remaining problem in the model setup is that in practice, only *R*_tot,*j*_, the total receptor expressions of each kind of a cell, can be experimentally measured, while the amount of free receptors at equilibrium, *R*_eq,*j*_, though being used extensively, is unknown. To find *R*_eq,*j*_, we first need to derive the amount of bound receptors of each kind, *R*_bound,*j*_, then use conservation of mass to solve *R*_eq,*j*_ numerically.

To calculate the amount of bound ligand *R*_bound,*n*_, we can simply add up all entries in the *n*-th column for every **q**’s:

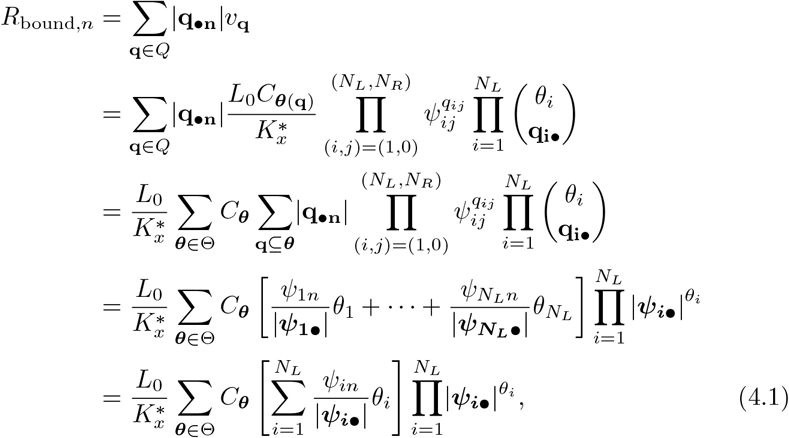

where 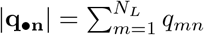, and 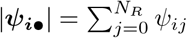.

By the conservation of mass, we have

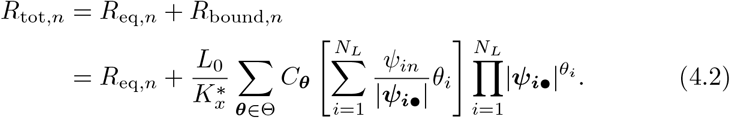

In this equation, *R*_tot,*n*_ are known, and any |***ψ***_***i•***_| is a function of every *R*_eq,*j*_, *j* = 1, 2,, *N*_*R*_, so all *R*_eq,*j*_ need to be solved together. This system of equations usually does not have a closed form and must be solved numerically. When implementing, we suggest taking the logarithm of both sides of these equations so the exponents can be eliminated and the range is restricted to positive numbers.

As a side note, the total amount of bound receptors regardless of kind is

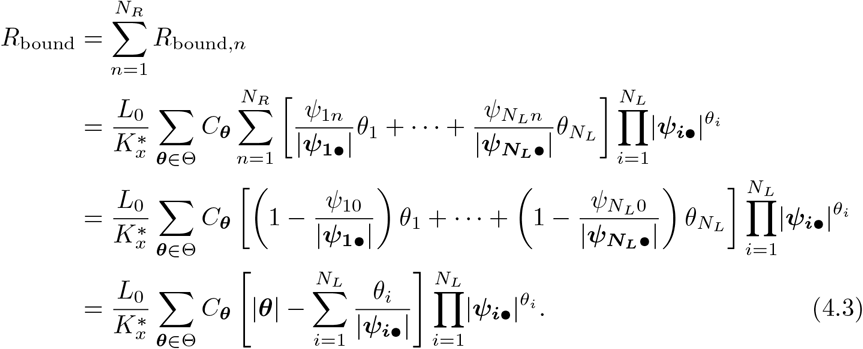

#### The amount of bound ligand complexes

Our model makes many macroscopic predictions readily accessible. For example, the amount of ligand bound at equilibrium is a useful quantity when measuring the overall quantity of tagged ligand. To compute this number, we can add up all *v*_**q**_ except the **q**’s that only have nonzero values on the 0-th column, *v*_0,eq_. Consequently, the model prediction of bound ligand at equilibrium is

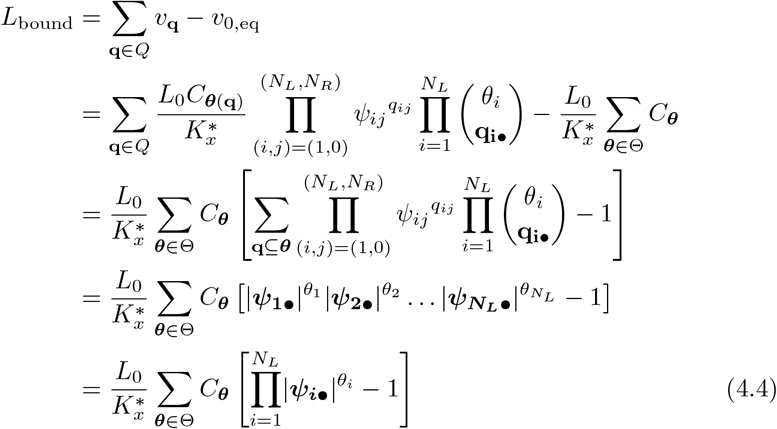

when 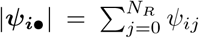, and the predicted amount of bound complex ***θ*** (complex of each kind) is

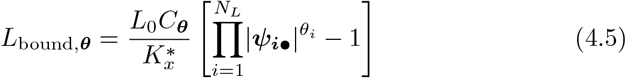

#### The amount of fully bound ligands

In multivalent complexes like bispecific antibodies, drug activity may require that all subunits be bound to their respective targets [11]. The predicted amount of ligand full-valently bound can be calculated as

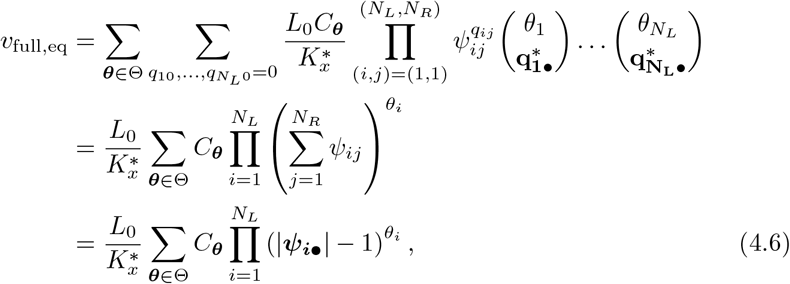

with 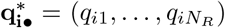, the **q**_**i**•_ vector without *q*_*i*0_. In this equation, the multinomial coefficient 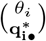 describes the number of ways one can allocate *θ*_*i*_ receptors to any position in the *i*-th row of the (*q*_*ij*_) matrix except the 0-th row which stands for unbound.

In fact, the predicted amount of any specific-valently bound ligands can be derived in such manner. For example, the amount of ligands that bind monovalently can be calculated as

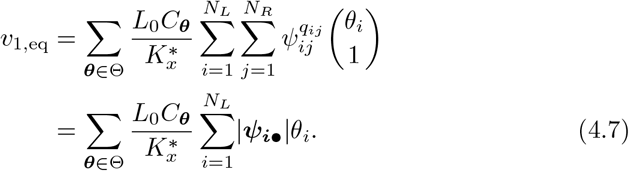

This can be used for estimating the amount of multimerized ligands, *L*_multi_ = *L*_bound_ − *v*_1,eq_, and multimerized receptors, *R*_multi_ = *R*_bound_ − *v*_1,eq_.

### 4.2. Complexes formed through random assortment

Another common mode of forming multivalent complexes in biology, such as in the formation of antibody-antigen complexes [4], is the stochastic assembly of monomer units to a common scaffold. Instead of a specific arrangement, we provide binding compounds of a fixed valency *f* and a mixture of monomer ligands, and complexes can form through random assortment. The concentration of these complexes, therefore, will follow a multinomial distribution.

To formulate this mathematically, we denote the proportion of *L*_*i*_ as *C*_*i*_, and 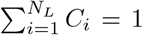. For example, say we have 40% *L*_1_ and 60% *L*_2_ in the solution to form dimers (*f* = 2), then *C*_1_ = 40%, *C*_2_ = 60%. As complex formation follows a binomial distribution, there will be 16% bivalent *L*_1_, 36% bivalent *L*_2_, and 48% *L*_1_−*L*_2_ complex. In general, the probability of complexes formed as described by ***θ*** in random assortment is

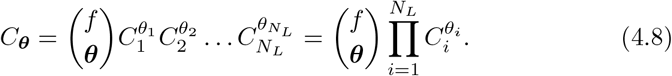

Since 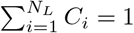, we know that

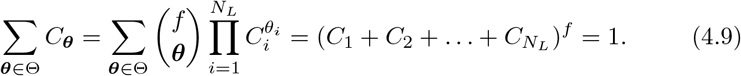

Plugging this relationship between *C*_***θ***_ and *C*_*i*_ into Equation 3.6 for the amount of a specific binding configuration, we have

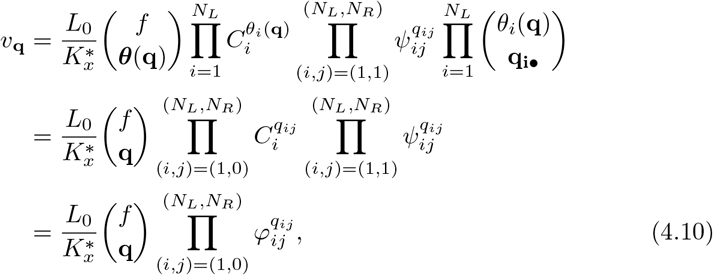

where 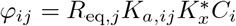 and *φ*_*i*0_ = *C*_*i*_.

#### Solve the amount of free receptors

Similar to the specific arrangement case, we still need to solve *R*_eq,*n*_ numerically from *R*_tot,*n*_. We first derive the amount of bound receptors of each kind at equilibrium as

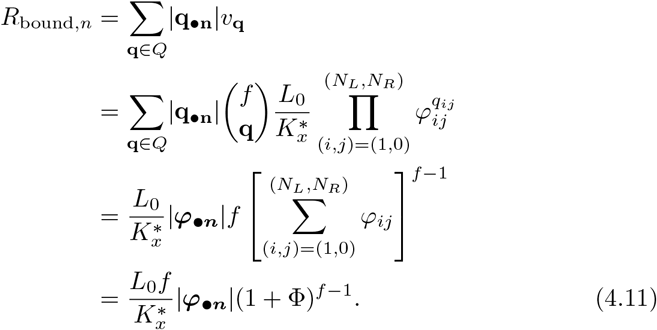

Then by the conservation of mass, we can numerically solve *R*_eq,*n*_ as

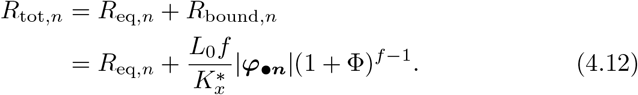

Again, since 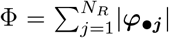 is a function of every *R*_eq,*n*_, all *R*_eq,*n*_ need to be solved together.

#### The amount of k-valently bound complexes

For randomly assorted complexes, we first derive the amount of ligands that bind *k*-valently. As we will show, it has a useful expression that can used to find many other quantities conveniently. First, let’s break **q** into two separate vectors, **q** = (**q**_•**0**_, **q**_•**x**_). We define the vector formed by the 0-th column of **q** which stand for unbound as **q**_•**0**_, and the one formed by the other elements as **q**_•**x**_. By the model setup, |**q**| = *f*, |**q**_•**x**_| = *k*, and |**q**_•**0**_| = *f* − *k*. We then have

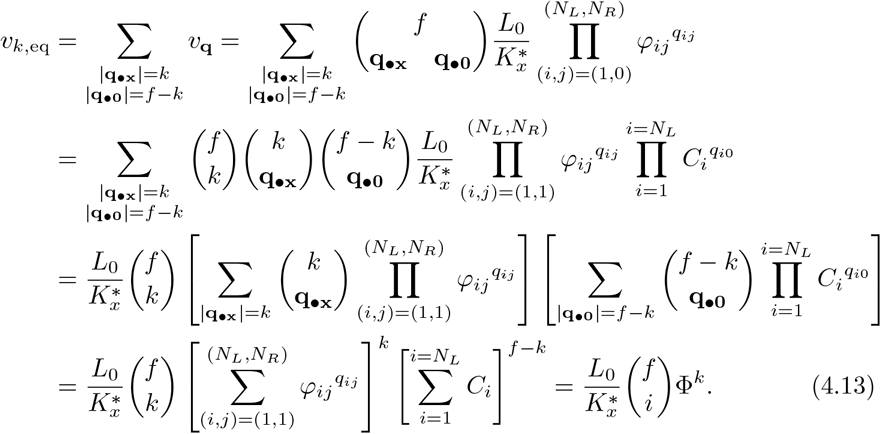

#### The amount of total bound ligands and receptors

Many macroscopic properties can be derived from *v*_*k*,eq_. For example, the amount of total bound ligands is simply the sum of ligands bound monovalently to fully, and can be simplified to

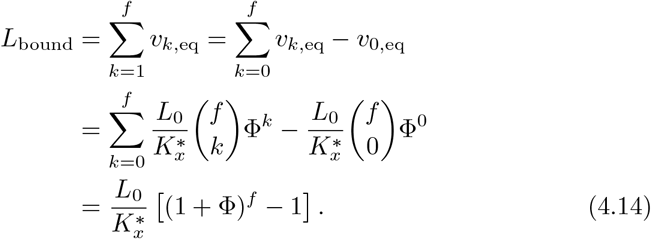

Similarly, the total bound receptors should be

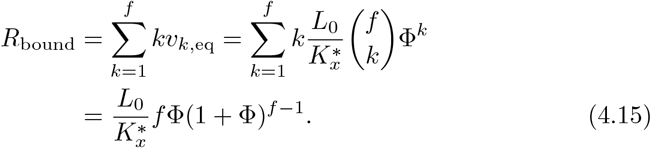

As we show here, these quantities all have elegant closed form solutions, and they are only dependent on Φ, a single value that incorporate all information about receptor amounts, monomer ligand compositions, and binding affinities. Φ was previously defined as 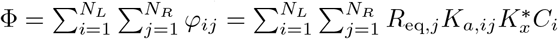.

#### The number of cross-linked receptors

In some biological contexts such as T cell receptor-MHC [3] or antibody-Fc receptor [4] interactions, signal transduction is driven by receptor cross-linking due to multivalent binding. The amount of total cross-linked receptors can be derived from *v*_*k*,eq_ as

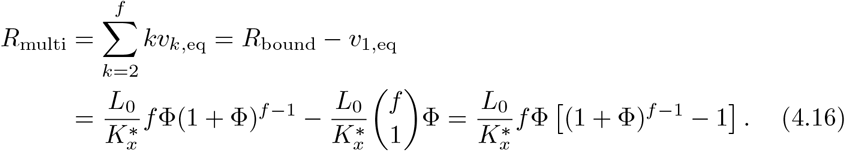

To find the number of crosslinked receptors of a specific kind, *R*_*n*_, requires extra consideration. Similar to how *v*_*k*,eq_ was found, we break **q** into three separate vectors, **q** = (**q**_•**0**_, **q**_•**n**_, **q**_•**x**_). **q**_•**0**_ is the vector formed by the 0-th column of **q, q**_•**n**_ is the vector formed by the *n*-th column of **q**, and **q**_•**x**_ contains all others. If the complex is *s*-valently bound, then |**q**_•**0**_| = *f* − *s*. We further assume that |**q**_•**n**_| = *t*, then |**q**_•**x**_| = *s* − *t*. By this setup, we have

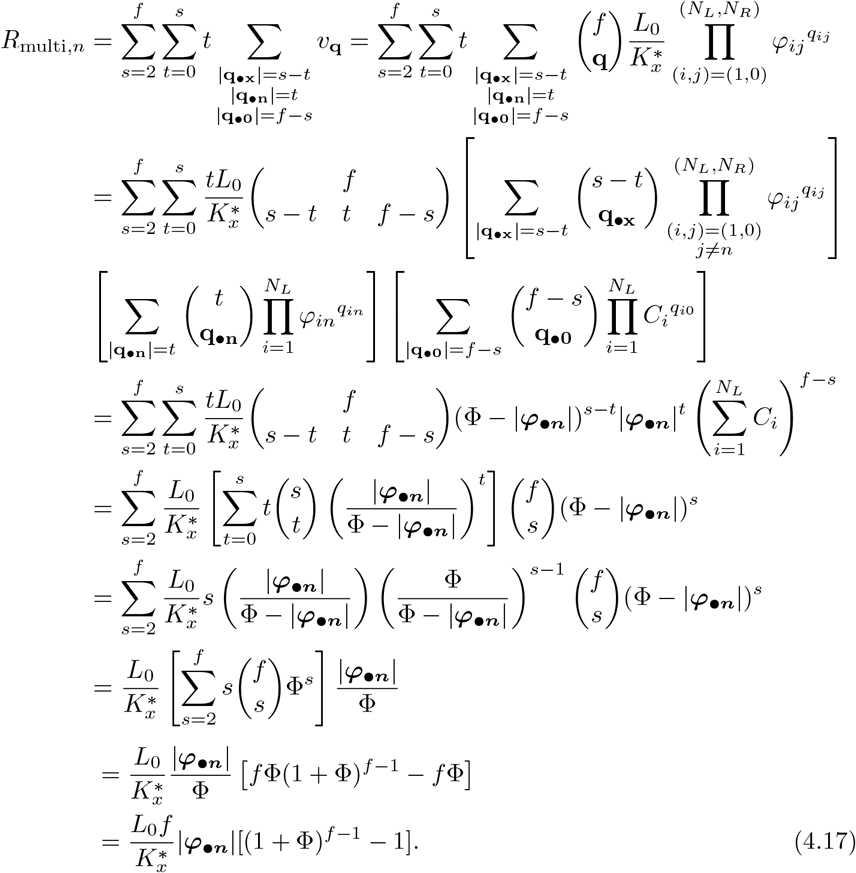

This formula can useful when investigating the role of each receptor in a pathway that requires multimerized binding.

Of course, the macroscopic predictions provided in this section cannot exhaust many biological quantities one may wish to study, but with the ideas demonstrated here, the readers can derive their own formulae as needed.

### 4.3. Numerical implementation notes

In this model, most predictions can be calculated directly by closed form formulae after *R*_eq,*n*_’s have been solved. For solving *R*_eq,*n*_’s numerically, we have not found issues with deriving numerical solutions generally, except at extreme affinities (e.g., less than 1 pM *K*_*d*_) combined with very high valency (e.g., greater than 64), where floating point errors can cause problems with the termination conditions of the root-finding operation. We have learned through experience that one need not set bounds on the root-finding, as the function is monotonic with a single root. While we use auto-differentiation of the model through the Python package `Jax` or Julia package `ForwardDiff.jl` during root-finding (both packages available on GitHub), one can use numerical differencing with identical results. Root-finding of *R*_eq,*n*_ is by far the slowest part of the calculations.

#### Ligand concentration handling

Our model requires the total concentration of ligands at equilibrium, *L*_0_. There are at least four approaches one could take to rectify some measurement of ligand concentration with the model: (1) First, one could apply an assumption of no ligand depletion. This is an extreme assumption in many cases but can be applicable in *in vitro* experiments where it is known that the ligand amount is many orders of magnitude greater than that of the receptors. (2) Alternatively, one might know the concentration of ligand in solution after some or all of any ligand depletion has occurred. (3) If one has an estimate of the absolute number of receptors and ligand units before binding, *L*_0_ can be solved numerically by conservation of mass along with *R*_eq,*n*_’s. We have

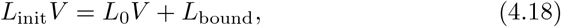

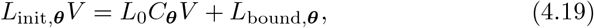

where *V* is the effective volume of the ligand solution, *L*_init_ is the total initial ligand concentration, and *L*_init,***θ***_ is the initial concentration of complex ***θ***. *L*_bound_ and *L*_bound,***θ***_ may be found at Equations 4.14 and 4.5, depending on the occasion. (4) Finally, certain aspects of the system may not be sensitive to ligand concentration as an input parameter, or one could treat concentration as an unknown.

## 5. Application examples

In previous sections, we have shown how all macroscopic predictions made in this work can be written in closed form formulae. Therefore, some computationally expensive analyses are enabled by the efficiency of our model. Here, we provide two examples to demonstrate the utility of large-scale predictions made possible by this model.

### 5.1. Mixture binding prediction

Leveraging a synergistic effect among two or more drugs is of great interest in pharmaceutical development. A challenge in investigating synergy is to identify its underlying source. Most biological pathways follow a similar pattern: when the drug binds to certain surface receptors of a cell, a downstream pathway in the cell is initiated, leading to some actions. Therefore in general, synergism can come from either the initial binding events themselves or downstream signal transduction interactions. Binding-level synergy means that merely using a combination of ligands boosts the amount of binding to the important receptors and thus intensifies the overall effect. Downstream effect synergy indicates that the benefit of using mixtures arises from other cellular regulatory mechanisms two ligands can bring about. The binding model we propose here can help to investigate this issue by offering accurate predictions for the binding of multivalent complex mixtures.

In Figure 3, we provide an example of mixture binding predictions (Figure 3a). We investigate a mixture of three types of ligand complexes, bivalent *L*_1_ (***θ***_**1**_ = [2, 0]), bispecific *L*_1_−*L*_2_ (***θ***_**2**_ = [1, 1]), and monovalent *L*_1_ (***θ***_**3**_ = [1, 0]). The crosslinking constant is set to be 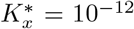 cell. M, similar to previous results [4]. We predict the amount of binding of this mixture to a cell expressing three types of receptors, with ***R***_**tot**_ = [2.5 × 10^4^, 3 × 10^4^, 2 × 10^3^] cell^−1^. The affinity constants of *L*_1_ to these three receptors are ***K***_***a***,**1•**_ = [1 × 10^8^, 1 × 10^5^, 6 × 10^5^] M^−1^, and of *L*_2_, ***K***_***a***,**2•**_ = [3 × 10^5^, 1 × 10^7^, 1 × 10^6^] M^−1^. Here, we investigate the changing concentration of ***θ***_**1**_ and ***θ***_**2**_, while holding the amount of ***θ***_**3**_ constant at 0.2 nM. Figure 3 shows the predicted total ligand bound (Figure 3b) and *R*_3_ bound (Figure 3c) for only ***θ***_**1**_ or ***θ***_**2**_ with concentration from 0 to 0.8 nM, and their mixtures in every possible composition with total concentration 0.8 nM.

**Figure 3:**
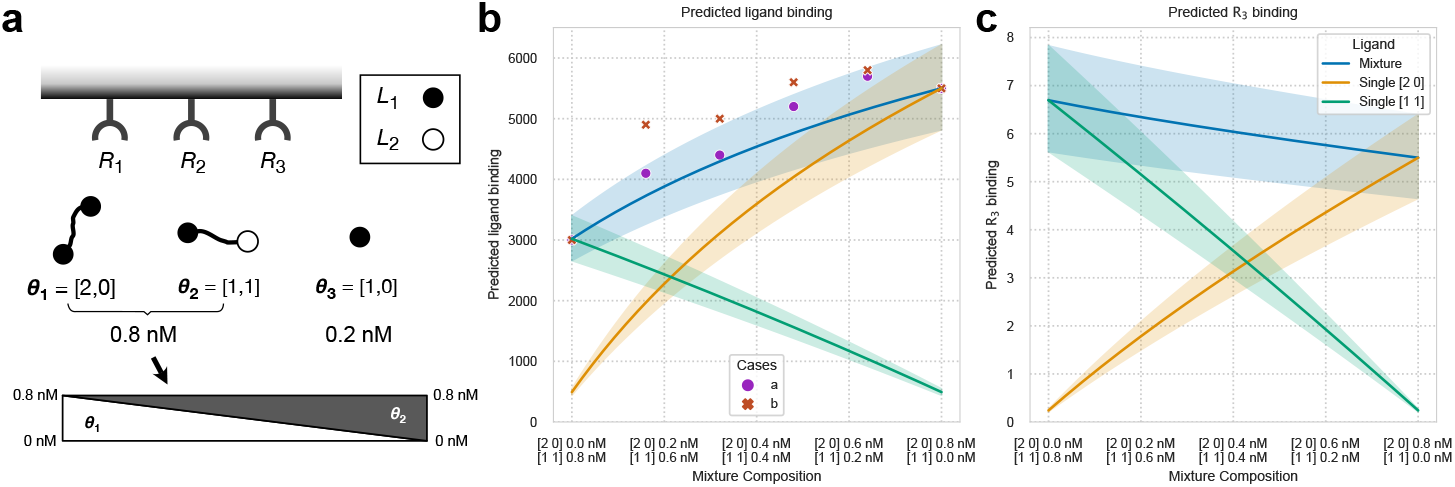
Prediction on mixture binding of ***θ***_**1**_ = [2, 0], ***θ***_**2**_ = [1, 1], and ***θ***_**3**_ = [1, 0]. (a) A schematic of the binding scenario; (b) The predicted total ligand binding; (c) The amount of bound *R*_3_ at equilibrium. Shaded areas in (b), (c) are simulated confidence intervals by varying the receptor levels up and down by 10%. The points on (b) represent possible experimental results, arbitrarily drawn for demonstration. In case *a* (purple circles), since most data points are inside the confidence interval, we can assume the measurement error can explain these variations. In case *b* (orange crosses), however, the synergism of these complexes are beyond the binding level.

Mixture binding prediction can help us identify the source of synergy. To connect model predictions to experimental measurements, ligand binding might be measured by fluorescently-tagged ligands, while the number of bound receptors of a specific type might associate with an indirect measurement such as cellular response. After making a series of measurements for different compositions of mixtures, we can fit the 100% of one complex cases (numbers on the two ends on the plot) first and then compare the mixture measurements to the predictions. Determining whether the downstream effect contributes to the observed synergy (or antagonism) can be framed as a hypothesis testing problem:

#### H_0_: The synergism of the mixture can be explained solely by binding

The uncertainty of mixture binding prediction comes from measurement errors of receptor abundance and binding affinities. Usually, the receptor expression of a cell population has an empirical distribution which can be measured. The confidence intervals in Figure 3 were drawn with the assumption that receptor expression fluctuates up and down by 10% (coinciding with log-normally distributed amounts). Also, due to the measurement technique, the binding affinities may be over- or underestimated [12]. The confidence interval of mixture prediction can be determined by the model with all these considered, and a *p*-value can be derived.

We arbitrarily drew some possible experimental results on Figure 3b for demonstration. If most mixture measurements fall within the confidence interval of the predictions (such as case *a* annotated by the purple circles in Figure 3b), the synergy will very likely come from binding only. However, if the measurements are obviously beyond the confidence interval (case *b*, the orange crosses), it is reasonable to suspect a synergistic (or antagonistic) effect beyond binding alone. With the flexibility of the binding model, this method can also be extended to a mixture of more than two compounds.

### 5.2. Binding space of a ligand

When a dose of ligand (drug, hormone, cytokine, etc.) is released into the circulation system of an individual due to either physiological response or exogenous administration, the compound will spread and bind to many cell populations to varying extents. An essential question in pharmacology is how much a compound will bind to their intended target populations compared to off-target ones. This question is important for understanding basic biology as well as developing new therapeutics. For example, hormones and cytokines are important signaling molecules, and having a quantitative prediction of on- and off-target binding can help us understand their mechanism greatly. For drug development, binding prediction can guide optimization to improve specificity toward the intended targets [13]. A cell population can be defined by the receptors they express. Therefore, given the parameters of the dose and the receptor profile of each cell population, our model can make all the predictions discussed previously.

From the perspective of this binding model, there is nothing special about any specific cell population. If the local concentration is constant everywhere, our model can map any cell with a certain receptor expression to the amount of binding induced by this dose. If the biological activity of this compound on a cell is related to the quantity of binding to a certain ligand or receptor, the effect of this dose can be written as a function *f*, with

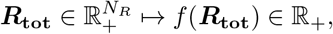

where ***R***_**tot**_ is a vector of nonnegative entries that describes the cell’s expression of *N*_*R*_ receptors, and *f* (***R***_**tot**_) is the amount of binding. Here, we define the binding behavior of this dose (or any compound) as its binding space.

In Figure 4, we plot the binding space of a bivalent *L*_1_ ligand ***θ*** = [2, 0] with concentration 1 nM. The binding affinities are the same as described in the last subsection (5.1). In this binding space, we consider three receptors, *R*_1_, *R*_2_, and *R*_3_. We plot how the amount of binding relates to the cell expression profile, ***R***_**tot**_. Here, the amount of *R*_1_ and *R*_2_ varies with the two axes, while *R*_3_ is held constant at 2.0 × 10^3^ cell^−1^. In this plot, each point represents a cell with a distinct expression profile, as some examples drawn on Figure 4a. Then we use colors and contour lines to show the amount of binding. From these two plots, we can see that although both ligand binding and *R*_2_ binding increase with more receptors, ligand binding is more sensitive to *R*_1_ amounts, and *R*_2_ binding *R*_2_ amounts. To consider any specific cell population, one only needs to determine where its expression profile falls on the plot and read the predictions from the contour line. For example, on Figure 4b, the red cell population will have about *e*^5.2^ = 181 bound ligands per cell. The number of contour lines a population ride on can also show intrapopulation variation. In this case, we expect the variation in ligand binding to fall between *e*^4.3^ = 74 and *e*^6.0^ = 403.

**Figure 4:**
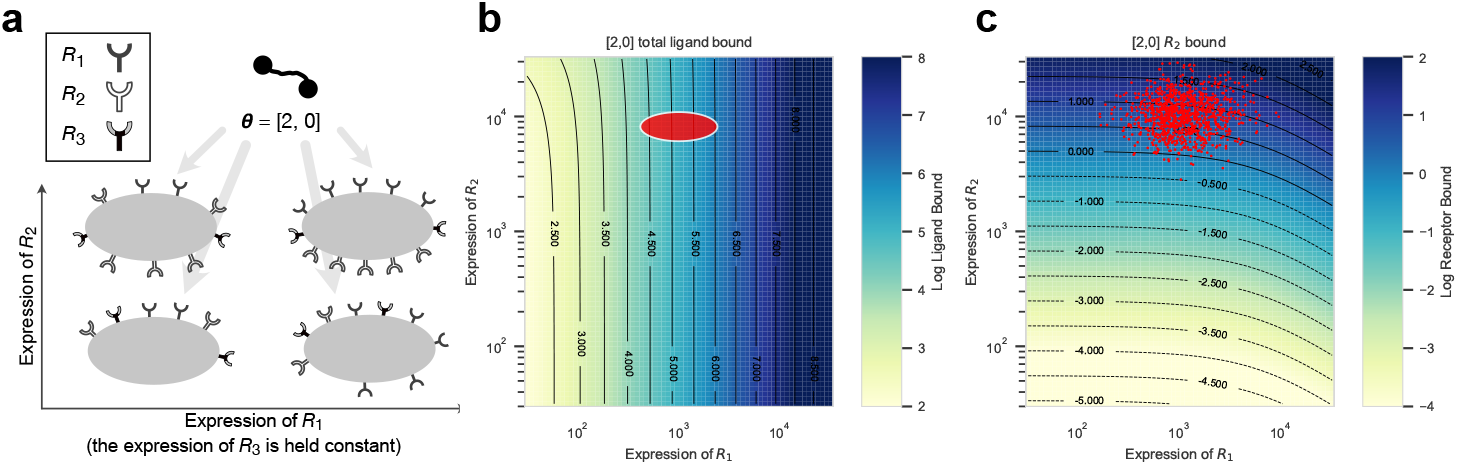
The binding space of 1 nM ***θ*** = [2, 0]. (a) A schematic diagram of the ligand and four examples of receptor-expressing cells represented by the coordinate; (b) The amount of total ligand bound; (c) receptor *R*_2_ bound predictions. The x- and y-axis show the expression of *R*_1_ and *R*_2_, while the expression of *R*_3_ is a constant, 2.0 × 10^3^ cell^−1^, and not shown. Any cell population can be drawn on the binding space. For example, the red ellipse on (b) represents a cell population with receptor expression of roughly ***R***_**tot**_ = [1.0, 10.0, 2.0] × 10^3^ cell^−1^. We can alternatively project points of experimental single cell expression data onto a binding space, as shown on (c) (the points were generated arbitrarily assuming a population of log-normally distributed *R*_1_ and *R*_2_ expression for demonstration purpose.

The binding space can provide ample information about the compound. It is an intrinsic property of a ligand given its concentration and other ligand it mixes with, independent of any specific cell. The biological process of drug diffusion to a certain cell is analogous to sampling a point from this binding space. Its gradient indicates in which direction the binding level increases the fastest, as well as to which receptor amount it is the most sensitive. An inactive antagonist that introduces binding competition with the ligand can distort its binding space, and we can visualize it by the change of shape in the contour lines. This plot can also intuitively demonstrate intrapopulation binding variance and interpopulation cell specificity of the compound. With the development of high-throughput single-cell methods such as flow cytometry, the expression profiles of a collection of cells can be identified en masse, and we can overlap their results onto a binding space plot (as in Figure 4c). This shows the promise of applying our model to single-cell data. Although we can only visualize two receptors in a plot, binding space applies to any *N*_*R*_ types of receptors. Theoretically, the concept of the binding space of a ligand is only complete when all relevant surface receptors are considered.

## 6. Discussion

In this work, we propose a mechanistic multivalent binding model that accounts for the interaction among multiple receptors and a mixture of ligand complexes formed by binding monomers. The flexible framework allows a mixture of both homogeneous and heterogeneous ligand complexes, even when they don’t have the same valency. We first derive the amount of ligand of a specific binding configuration at equilibrium through the law of mass action. Using this formula, we make macroscopic predictions by applying the multinomial theorem strategically. Our predictions cover cases where complexes are formed by specific arrangement or random assortment. Finally, we provide two practical examples of how this model can help with biological research.

Compared with previous approaches, the model here is a uniquely scalable and elegant approach to multivalent binding when considering multivalent complexes of heterogeneous monomer composition and/or multiple receptors. Scalability to higher valency complexes is essential as rule-based computational models become impractical due to a combinatorial explosion of binding states. By contrast, our model can make a large number of predictions easily, enabling mixture synergy analysis and binding space calculations across individual cells.

The mathematical elegance of the model welcomes analytical studies and incorporating it into higher-level computational frameworks. For example, we apply auto-differentiation to ensure accuracy in the root-finding operation when solving for unbound receptor. We have similarly used auto-differentiation to solve for the gradients of the model with respect to input quantities when fitting it to data points. One could even feasibly derive analytical forms of the gradients. This enables one to build more complex computational models on top of this binding framework, such as inferring the composition of multivalent complexes in solution from indirect high-throughput assays. While differentiation of differential equation models is possible through adjoint state methods, solving can be sensitive to the parameters of the system, is much less efficient, and requires trade-offs in accuracy for performance.

The assumptions made in this model may compromise its accuracy in some cases. Our setup has a single crosslinking constant, 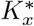,to reflect the multivalency effect. In practice, this model has worked well in predicting experimental binding results [13, 14, 4]. However, the steric effects of a multivalent ligand can be more complicated and context-dependent. The complication of multivalency effect comes from the geometry of ligand complexes that introduced steric effect as well as the distribution of receptors on the cell surface. For instance, the length of the hinge region is needed to estimate the radius of area a molecule can reach [15]. Receptor clustering can be play a big role in the behavior of ligand binding as well [16]. Accounting for these effects requires more in-depth studies than just measuring the monomer binding kinetics. Some other computational approaches investigate steric effects more meticulously, but inevitably introduce some added complexity. For example, previous work has conducted a case-by-case exploration of how ligands bind when distributed randomly or ordered, arranged as a lattice, ring, or chain to give a better hindrance factor estimation [10]. When the actual situation is not known, our model can serve as an adequate starting point.

Although this model is very general purpose, it mainly focuses on the binding dynamics on a cell surface, similar to the previous work on which it is based [7, 8, 5]. For ligands discordant with the two-step binding process shown in Figure 2, other model constructions might be necessary. For example, some previous work focuses on scaffold proteins as intracellular multivalent complexes [17], but these often lack independence between the individual monomer binding events. In this case, various alternative computational models have been developed [18, 19, 20].

Surface receptor binding is a universal event in biology. A prevalent question calls for a general solution. We expect this model to be successfully applied to many contexts. Previously, we have used a simpler version of the random assortment model to accurately predict IgG antibody-FcγR interactions [4], and also applied it to fit epithelial cell adhesion molecule binding data [13, 14]. We are also working on applying the model to IL-2 immunocomplexes [21], for optimization of high-valency cytokines with specific cell targeting [22], design of cytokine-antibody bispecific antibody fusions, and as a factorization kernel in dimensionality reduction of systems serology data [23]. With the arise of multispecific drugs in the recent decade [24], we expect this model to apply even more widely, exhibit its full competence and facilitate both basic scientific research and new therapy development.

## Author contributions

Z.C.T.: Methodology, Writing – original draft; A.S.M.: Funding acquisition, Writing – review & editing.

## Data and software availability

A Python package of this model and the code for the plots can be found at https://github.com/meyer-lab/valentBind/. We also provide a Julia package of the model at https://github.com/meyer-lab/polyBindingModel.jl/.

